# The role of resource defensibility in facilitating sexually-selected weapon evolution: An experimental evolution test

**DOI:** 10.1101/2024.12.06.627045

**Authors:** Tyler Audet, Audrey Wilson, Reuven Dukas, Ian Dworkin

## Abstract

Animal weapons have evolved multiple times primarily for battling for access to mates. Despite intra-sexual selection being common, weapon evolution has evolved relatively rarely. So why do weapons not evolve more commonly? It has been hypothesized that three precursors are necessary for the initiation of weapon evolution: high variance in reproductive success, patchy high-value resources, and spatial environments conducive to one-on-one competition. Here, we test this hypothesis by performing experimental evolution in *Drosophila melanogaster*, utilizing heterogeneous environments where conditions facilitating territorial defense and opportunities for competitive interactions vary. We examine changes in sexually dimorphic morphology and male aggression that are predicted to occur, based on this model. We also examine whether condition dependence for sexual dimorphism has evolved after 35 and 75 generations of experimental evolution. Aggression did increase, albeit modestly, in environments that facilitate resource defense. Morphological changes are modest although with some trait specific changes to allometry, generally in the opposite direction of our predictions. Condition dependence trends in the opposite direction from those predicted by our hypothesis as well. We discuss our results in the context of the necessary conditions for the evolution of weapons, and if, and when condition dependence of trait exaggeration may evolve.

## INTRODUCTION

The expression of exaggerated sex-specific traits in the animal kingdom has long been an area of fascination with biologists (Darwin 1871). These elaborate traits evolve along a spectrum, where ornaments, often presumed to function in signaling during inter-sex interactions, lie on one end, while weapons used intra-sexually are on the other, although with many traits serving both functions (Kokko et al. 2002). Within weapons, a continuum also exists, as direct combat can result in damage to individuals who engage in it. Although weapons may be used for physical combat, many ‘weapon’ traits are used for agonistic inter-sexual signalling, rarely resulting in physical escalation; such as the stalk-eyes of Diopsid flies (reviewed in McCullough et al. 2016). More generally, weapon size may serve as a general cue to conspecifics with regards to potential outcomes in direct combat. In *Drosophila melanogaster* agonistic threat displays and “escalated” physical altercations, are genetically correlated (Baxter et al. 2023). It has long been understood that traits evolve along both the continuum, with respect to the relative contributions of intra- and inter-sexual selection in general, and for the continuum for physical vs. signaling/cues for intra-sexual agonistic interactions specifically (Andersson 1994).

The intensity of sexual competition has for a long time been attributed to the accessibility of a limiting sex (typically females) by a less limiting sex (typically males; Darwin 1871; Bateman 1948). In large part due to the consequences of anisogamy, including female receptivity and parental investment, the operational sex ratio (OSR) of populations can often be male biased. This male biased OSR leads to various forms of competition for access to mates including sperm competition, mate harm, and intra-sexual aggression, which can result in a high variance in the number of matings individual males acquire relative to females in the population (Nandy et al. 2013; Bath et al. 2021; Sepil et al. 2022). The ability to differentially access mates can become more intense depending on the mating system of the organism in question, as well as ecological constraints (Emlen and Oring 1977). The Environmental Potential for Polygamy (EPP) has been hypothesized to depend on the ability to defend multiple mating partners, or resources desired by multiple mates, by an individual (Emlen and Oring 1977). It also depends on the ability of an individual to allocate sufficient time defending this resource, without the requirement of pursuing other activities such as parental care (Emlen and Oring 1977). The prevalence of sexual selection and EPP may be quite common in the animal kingdom, with competition over mates occurring in many species. However, the evolution of weapons is far rarer than the occurrence of male-male competition for mating (Voje 2016; Palaoro and Peixoto 2022).

Traits under stronger sexual selection tend to scale disproportionately with body size compared to non-sexually selected traits, at least in some taxa (Simmons and Tomkins 1996). Traits under direct sexual selection may be used as indicators of size, either through direct female selection (Summers and Ord 2022) or male-male competition (McCullough et al. 2016). In the case of male-male competition for access to mates, larger traits can increase the chance of winning duels, and thus increase resource holding potential and potentially unequal access to mates (Palaoro and Peixoto 2022). While it has been proposed that there may be differences in patterns of “weapon” allometries (i.e proportional changes in weapon trait size with proportional changes in body size), depending on whether the trait functions directly in physical interactions between competing males, or provides threat signal or cue to assess rivals size (Eberhard et al. 2018; McCullough and O’Brien 2022). However despite this potential separation of roles (physical weapons vs threat signals), these structures generally show positive allometries, and are often hyper-allometric (proportional increases in weapon size is greater per unit increase in size), function along a continuum of roles (physical altercations through signaling), with a genetic correlation among the forms of aggression (Kodric-Brown et al. 2006; Voje 2016; Emberts et al. 2021; McCullough and O’Brien 2022; Baxter et al. 2023).

Static allometry is the log-log regression of trait size on body size, for individuals of a particular developmental stage. With regards to weapon evolution, this is for sexually mature adults, and may influence outcomes in male-male agonistic signaling or combat (Voje 2016; Eberhard et al. 2018). Allometric slopes of ∼1 (isometry) occur when increases in (log) trait size and (log) body size increase at a similar rate. Slopes below 1 (hypo-allometry) indicate relatively smaller trait values for a given body size (as body size increases). Slopes above 1 (hyper-allometry) denote disproportionally large trait values relative to body size (Bonduriansky and Day 2003; Voje 2016). Weapons, broadly defined, show a range of allometric coefficients, although they tend towards hyper-allometry across many taxa (Voje 2016; Eberhard et al. 2018; McCullough and O’Brien 2022).

In mating systems where EPP is high, and males compete for access to mates, intra-sexual selection may be intense and resource defense polygyny may create a skew in which males must fight for access to mates, favoring aggressive interactions. If the resource is sufficiently high value, males with high resource holding potential (RHP) should be more willing to escalate agonistic interactions (Hurd 2006). In primates, aggression and physiological markers of aggression such as the size of brain regions associated with aggression, are correlated with higher degrees of sexual selection (Lindenfors et al. 2007). A pattern has also been present in the literature that species with weapons tend to be more aggressive than closely related species without, or even individuals of the same species without the trait exaggeration (Moczek and Emlen 2000; Kudo et al. 2017; Boisseau et al. 2020). To date, to the best of our knowledge, the evolution of aggression in relation to the evolution of trait exaggeration has yet to be studied. It is known that males without obvious trait exaggeration, such as the pomace fly, *Drosophila melanogaster*, can be aggressive and show resource defense/territoriality (Dow and Schilcher 1975; Hoffmann and Cacoyianni 1990; Chen et al. 2002; Guo and Dukas 2020). This includes both the use of threatening signals, such aswing displays, and physical altercations mediated by use of the front legs and heads (details discussed in methods). *Drosophila melanogaster* selected for increased territoriality, showed increased mating success and longevity under some conditions (Hoffmann and Cacoyianni 1989). Other *Drosophila spp*, such as Hawaiian *Drosophila* show trait exaggeration (Spieth 1981), in particular hypercephaly, lekking behaviour and substantial aggression (Kudo et al. 2017). In the stalk-eyed *D. heteroneura*, head width is a strong predictor of male-male contest outcomes, as well as influencing mating success with females (Boake et al. 1997). One species of the melanogaster spp group, *D. prolongata* (part of the rhopoloa clade), has exaggerated male forelegs used in male-male combat, and outcomes of contests influence mating success (Toyoshima and Matsuo 2023). Although the native mating substrates for *D. prolongata* are currently unknown, the intensity of aggression in this species suggests that aggressive interactions are a necessary precursor for changes in male-biased sexual dimorphism. Increased competition for mates and associated increases in variance in reproductive success may lead to the evolution of increased aggression and contests, and weapon evolution.

Despite the prevalence of males engaging in intra-sexual agonistic interactions that contribute to mating success the relatively rare evolution of exaggerated weapons, begs the question as to why, when, and how weapons evolve. Building on the foundations laid out in Emlen and Oring (1977), Emlen (2008, 2014) hypothesized that three explicit conditions are necessary for the precursors of weapon evolution. First, there must be competition for access to females, likely in a way that creates asymmetry in access to mates, generating increased variance in male reproductive success. This may result in males who expend resources into trait (weapon) expression, as investing in increased resource holding potential may be crucial for reproductive success. Second, there are limiting, localized (patchy) resources required by females. If resources are distributed abundantly throughout the environment, there is little benefit in defending one patch if there is a plethora nearby of equal value that females may visit instead. Discrete patches of limiting resources result in predictable locations that females must visit, and therefore specific locations to defend. Finally, the layout of these resources must be such that males compete in duels, or one-on-one fights. If resource patches are sufficiently large (spatially) resource defense may be impossible, as many males attempt to usurp the dominant male at the same time. In this scenario, competition becomes a scramble and there is likely no direct benefit in being the strongest because a scramble does not necessarily reward the largest weapon, but rather the fastest male to secure a mating. These conditions correlate with observations of extant species with weapons (Emlen and Philips 2006). Dung beetles largely take up two strategies to sequester dung for their larvae; they may roll dung away from the source and bury it elsewhere, or they may dig tunnels adjacent to the dung source. If dung is rolled away, males scramble to fight over the dung ball largely out in the open, and many males may compete at once. Tunnels, however, restrict access to dung in a way where males interact in one-on-one competition for access to the female who requires the dung for egg laying. Phylogenetically, only in lineages where males interact in these restricted spaces allowing one-on-one duels have horns evolved, and they have never evolved when competition occurs as a scramble (Emlen and Philips 2006). Diospid flies also have species with exaggerated male eye-stalks used in aggressive signaling as well as species with only rudimentary eyestalks. Sexually dimorphic species form nocturnal clusters on rootlets where males are able to control access to multiple females, and due to the linear nature of the rootlets, interactions between males occur one-on-one with the larger male typically winning (Wilkinson and Dodson 1997). The monomorphic species appear to not display the same clustering behaviour that allows male-male competition for access (Wilkinson and Dodson 1997). Although the conditions laid out by Emlen (2014) correspond to weapon evolution in some taxa, little has been done to experimentally test if these ecological conditions are necessary and/or sufficient to initiate weapon evolution, or contribute to increased male-biased sexual size dimorphism.

During the bout of strong directional selection occurring during exaggerated trait evolution, rapid depletion of genetic variation could occur, as alleles influencing trait size fix in the population (Taylor and Williams 1982). It has been hypothesized that alleles for sexually selected traits co-opt alleles for condition, resolving this ‘lek paradox’ through this process of genic capture (Rowe and Houle 1996). The association between trait and condition results in a heightened condition dependent response to environmental perturbation, meaning that when conditions are poor the trait reduces in size to a greater degree than other non-secondary sexual traits (Rowe and Houle 1996). This disproportionate decrease in trait size creates an honest signal to potential mates, or opponents. This patterns has been observed in nearly every system where it has been studied (David et al. 2000; Bonduriansky 2007; Johns et al. 2014), with few exceptions (Although see: Fairbairn 2005; Ceballos and Valenzuela 2011; Perdigón Ferreira et al. 2023). For this reason, it could be anticipated that the evolution of condition dependence co-evolves with trait exaggeration. Whether this occurs early or late during exaggerated trait evolution is unknown, and it has been suggested that traits that are already more condition dependent may be co-opted first, and exaggeration occurs as a consequence (Johnstone et al. 2009).

To test Emlen’s (2008; 2014) hypothesis for the ecological conditions necessary for weapon evolution, we performed experimental evolution using *D. melanogaster*. We generated three experimental environments, including two conditions conducive for males to attempt to defend food resource desired by females (food optimized to maximize female fecundity). These defensible resources should result in male-male competition for monopolization of the resource and an increase in mating success of the dominant males. Size of resource patches was based on prior research that demonstrated that when resource access (via food patch size) was limited, male *D. melanogaster* would increasingly perform resource defense (Hoffmann and Cacoyianni 1990). We generated three environments where individuals had spatially constrained access to resource patches, potentially facilitating increased one-on-one contests in males. One where patch size was large and easily accessible, one where patches occurred in sizes conducive to resource defense attempts, and one facilitating the opportunity for one-on-one competition, via restricted openings leading to resource (Figure 1, Figure S1). Previous work demonstrated that depending on the spatial constraint of resources and opportunities for sexual selection, there was variation in efficacy of selection to purge deleterious alleles (Wilson et al. 2021). Based on the hypothesis set out by Emlen (2008, 2014), and the fact that male *D. melanogaster* extensively use their forelegs (prothoracic) in agonistic interactions (Chen et al. 2002), we predicted evolution of increasingly male-biased SSD for these legs, and an evolutionary increase in the allometric slope of this leg (relative to overall body size).

**Figure 1:**
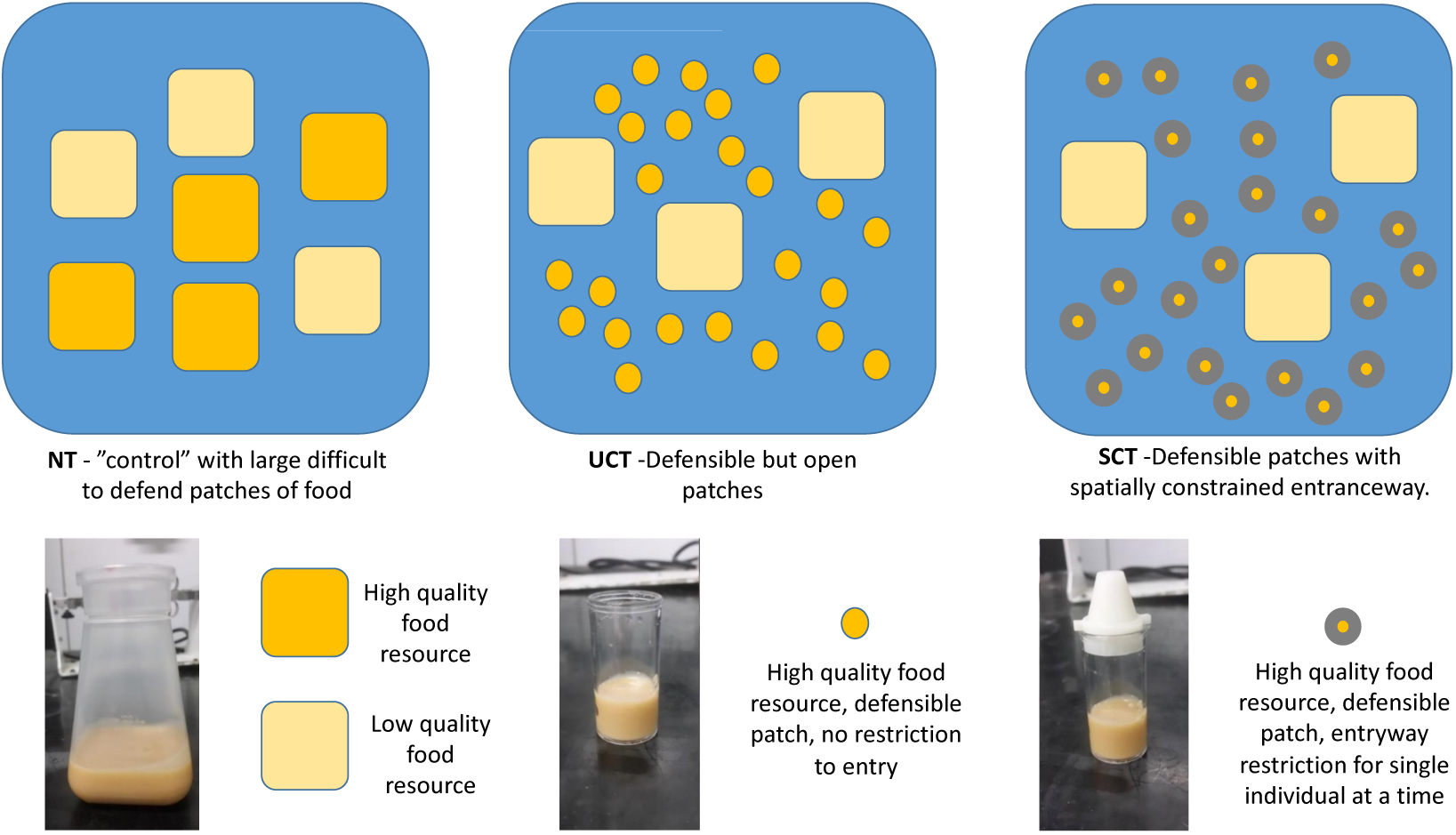
top-down layout of high- and low-quality food resource structure in each territorial treatment. Large squares represent open, easily accessible food, small circles represent food containers of a size *D. melanogaster* has been demonstrated to defend, and small circles with a grey ring represent resources with restricted access points. Darker objects represent high quality food and lighter objects represent diluted food. Placement of food containers in each cage, was random each generation. Lower row: side view shows the food containers and accessibility of resource. Smaller vials were decreased further in height after generation 10 to reduce the amount of space within the vial for flies to occupy (see methods).

Specifically, we predicted rank order changes based on three experimental environments we set up. We predicted a related response in wing length, as wings are used as a threat display. In the environment without small, defensible resource patches (large open resources, which single males cannot defend termed ‘no territory’, NT, hereafter), scramble competition tends to dominate (Hoffmann and Cacoyianni 1990). When resources are readily available in relatively large patches, scramble competition is classically believed to be the most common mating system in *Drosophila melanogaster* (Spieth 1974; Partridge et al. 1987; Hoffmann and Cacoyianni 1990) with recent work suggesting that interference competition may represent a substantial fraction of mating interactions (Baxter et al. 2018). We did not predict any substantial evolutionary changes in SSD or trait allometries in this environment, as a result of either of these mating strategies. In the second environment, with open, but defensible resource patches, termed ‘unconstrained territories’ (UCT), we predicted modest evolutionary increases in aggression, evolution towards male biased SSD and positive allometry in the legs. In the final environment, with spatially constrained access to resource patches, termed ‘spatially constrained territories’ (SCT), we predicted increased magnitude of evolutionary changes in aggression, SSD and positive allometry. We also evaluated traits for evidence of the evolution of increased condition dependence in male forelegs and wings relative to other traits. We predicted rank order differences in condition dependence across the three experimental environments (NT < UCT < SCT). While we observed modest changes in male aggression in the predicted direction, our results relating to changes in SSD, allometry and condition dependence were not generally consistent with our predictions. We discuss our findings in the context of the evolution of trait exaggeration.

## METHODS

### Environmental Treatments

Three environmental treatments were created to encapsulate the ideas of the three conditions for weapon evolution proposed by Emlen (2014). All three environments had the same surface area of high and low-quality food, and approximately the same volume of food. The “high quality” resources have been optimized for female fecundity based on nutritional geometry studies (Lee et al. 2008; Maklakov et al. 2008; Tatar 2011; Reddiex et al. 2013; Jensen et al. 2015), and were developed as highly desirable resources for female oviposition. Fecundity selection is an important driver of overall fitness in *D. melanogaster* (Tanaka and Yamazaki 1990), and evidence supports that it is likely one of the major contributors to the ancestral pattern of female biased SSD (Honěk 1993; Reeve and Fairbairn 1999). As such, female access to the high-quality food contributes to an individual’s ability to maximize fitness. “Low-quality” resources are dilutions of the high-quality resource, and for all three experimental environments, provided as four 177ml Drosophila culture bottles filled with ∼50ml of a 25% dilution of the high-quality resource. At generation 48, the low-quality resource was changed to a 10% dilution, as it was observed that flies were possibly beginning to adapt to this dilution, evidenced by a few normal sized larvae present in the diluted media. The purpose of these diluted food resources was to allow adult individuals to feed (even if the media could not generally support larval growth). In this way, individuals were not competing for resources necessary for adult survival, but for resources necessary to maximize reproduction. We note that we regularly observed individuals in each population using low quality resources for feeding, and some egg laying, but that larvae developing on them were small, developmentally delayed, and did not eclose as adults fast enough to contribute to the following discrete generation.

While all experimental environments shared the same amount (based on surface area) and proportions of high and low-quality food, they differed based on patchiness, accessibility and defensibility of resources (Figure 1). The “non-territory” (NT) treatment was designed to have large, open, easily accessible food patches that are difficult to defend as territories (Hoffmann and Cacoyianni 1990). NT effectively serves as a “control” environment, mimicking conditions of the lab domesticated population in spatial structure, and where scramble competition in *Drosophila* dominates (Hoffmann and Cacoyianni 1990). The “unconstrained territory” (UCT) treatment was designed to have small, defensible, but accessible patches. That is, while they are defensible, they are easy to access by multiple rival individuals, harder to monopolize, and potentially less likely to result in frequent duels. The third environment, the “spatially-constrained territory” (SCT) treatment, has the same patch sizes and numbers as UCT but access to the food patch is constrained via 3D printed “funnel caps” on each resource patch, restricting the opening (accessibility) to the resource (see Figure S1; and Wilson et al. 2021). These funnel caps are 3D printed plastic conical fittings that cover the opening of short vials, sitting on top with a bevelled edge leading to a smaller entrance at the peak with a 4mm opening. The addition of the funnel cap facilitates opportunities for defending and holding resources, and provides additional physical spaces that could encourage agonistic encounters for control of the resource to occur as one-on-one battles. The 4mm aperture is large enough that two males could pass each other, but small enough that one male can harass and attempt to restrict access to other males. Each environmental treatment was set up in mesh BugDorm-4F3030 cages (30cm) with the specific set up as follows; the NT treatment had four Drosophila culture bottles (177ml, 5.5cm length and width for a total surface area of 30.25cm for a total of 121cm total high quality food surface area) each containing ∼50ml of the high-quality food resource with four drops of a yeast and orange juice mixture placed on top to attract females (Dweck et al. 2013). The UCT treatment had 25 open vials (height of 32mm, 25mm outer diameter, ∼22mm inner diameter, 4.8 cm^2^ surface area for a total of 120cm of high quality food surface area), close to the optimal 20mm diameter that promotes resource-defence polygyny (Hoffmann and Cacoyianni 1990), each with a single drop of yeast-orange juice paste on the food surface. The SCT treatment had the same set-up as UCT (including surface area and volume of resources), except each vial had a 3D-printed funnel cap (22mm diameter, 25mm height, 4mm opening) to restrict access to the vial (resource patch) with a smaller entrance. Vial heights were reduced for UCT and SCT treatments from 95mm to 32mm at generation 10 of experimental evolution to reduce the amount of space between the top of the funnel caps and the surface of the resource in the SCT treatment. This was done to increase defensibility of the resources as initial monitoring of these vials showed high adult densities. Pipe cleaners were wrapped around the tops of bottles and vials containing high-quality food resources as perching sites. The outline of the territorial treatments and food containers can be seen in Figure 1 and Figure S1.

### Experimental Evolution Population Maintenance

Populations were created by collecting virgin females and males from a large outbred, lab-domesticated population, initiated from a large collection (several thousand individuals) from Fenn Valley Winery (FVW), Michigan in 2010 (GPS co-ordinates: 42.578919, −86.144936). This population had adapted to lab conditions for ∼160 generations, prior to the initiation to this experiment. Thus, confounding effects of concurrent selection for lab adaptation would be minimized (Harshman and Hoffmann 2000). From this population, 300 males and 300 females were placed into a cage, set up with one of the three environmental treatments. This was done with 4 replicate cages for each treatment, resulting in 12 lineages total (4 independent lineages per treatment). Populations were maintained at 12L:12D cycles at 21°C with 60% relative humidity in a Conviron walk-in chamber (CMP6050). The populations were kept on a 13-15 day schedule depending on emergence times, such that each population had about an equal amount of adults contributing to the next generation (census size was not measured directly). After the initial populations were placed into their respective treatments, adults were allowed to mate and lay eggs for three days. After this period, the media with eggs and larvae was removed from these cages and placed into new cages (without adults) to allow for development and eclosion. Development and eclosion occurred over a 10-12 day period. Once the new generation of adults emerged, old food was removed, and new food was placed in these cages with the set-up described above, and the cycle was repeated. This time-frame was used, as it was too short for the emergence of the rare individuals who developed on the low-quality resource (which had very few pupae regardless). For each generation, the new food was placed into the cages in a random distribution, and the cages were placed onto racks in a random order, such that each population varied in position in the walk-in chamber each generation.

### Assessment of male competitive fitness across the environmental treatments

To assess how potential spatial constraints and opportunities for territoriality interact to influence variance in male mating success, we performed an experiment to assess male competitive fitness. We predicted that males reared on high quality resources would have increased competitive fitness (compared with the common competitor) than those deprived of food during their terminal growth period (and are therefore smaller), and these differences would increase with opportunity for territoriality. For this experiment (summer 2020) we used the “ASW” population of *Drosophila melanogaster* established from 600 field collected females during the spring and summer of 2018 at various sites near Hamilton, Ontario, Canada, and maintained at census size above 2000 individuals each generation. For more details about this population, please see (Scott et al. 2022). The spontaneous X-linked c*rossveinless* (*cv*) mutation was introgressed into this population to serve as a visible marker to assess competitive fertilization. This visible mutation was chosen as previous work in the lab demonstrated that it had relatively modest deleterious effects in comparison to many visible markers. To manipulate male quality we placed 50 eggs onto high quality food and allowed high quality flies to develop to eclosion at 24 C before collecting virgin flies, while low quality males were removed from the food two days prior to pupation as described below. After eclosion, flies were stored in individual vials prior to the experimental manipulations. One individual focal ASW male of either high- or low-quality, and one *cv* male were then aspirated into a test cage (355ml plastic containers) with high quality resource corresponding to the treatments described above as well as an open low quality resource patch (high quality food diluted to 25%), sufficient for adult feeding and hydration, but insufficient to support proper growth. NT treatments contained a 1oz plastic cup with ∼35mm surface area of high-quality food, UCT contained a ∼22mm surface area vial of high quality food, and SCT contained the same food vial as the UCT treatment but with the restricted funnel cap with a 4mm opening as described above. Focal and *cv* males were allowed to settle into their environment for two hours before a virgin homozygous *crossveinless* female was introduced. Because of the X-linked recessive nature of *cv*, females sired by a *cv* male would be phenotypically crossveinless, and females sired by the focal ASW males would by phenotypically wild-type. Offspring from each treatment were allowed to develop and then collected for phenotyping to count the number of phenotypically crossveinless and wild-type female offspring.

### Condition Manipulation

At generation 35 and 75 of experimental evolution, two 177ml Drosophila food culture bottles containing high quality food were placed in each environmental treatment after the initial 3-day egg laying period for population maintenance. These bottles were removed after 7 hours to keep egg density low, and were kept at 21°C. Upon emergence of adults, 20-25 pairs were placed in 3 containers per replicate with a 2% apple-juice agar plate with a drop of orange juice yeast paste on the surface. Eggs were collected and placed into vials containing high-quality food resource at a density of 50 eggs per vial. For each of the 12 populations, 16 vials of eggs were collected and were split into three condition cohorts to undergo food deprivation protocol (Stillwell et al. 2011). The purpose of the food deprivation was to manipulate organismal condition, generating size differences, by limiting the nutritional content available to the larvae during growth phases of development. The first condition cohort (0) has normal food availability throughout larval development. Condition cohorts 2 and 1 each have successively increased days of food restriction before the end of larval development, with cohort 1 spending one day before the end of larval development without food, and cohort 2 with two days without food. Condition cohort 0 consisted of 4 vial replicates and developed on food for 6 days, cohort 1 consisted of 5 vial replicates and was left to develop on food for 5 days, and cohort 2 consisted of 7 vial replicates and was left to develop on food for 4 days. After these time periods the larvae from cohort 1 and 2 were removed by adding 5ml of a 40% sucrose solution to each vial and shaking for 20 minutes. Once the larvae were loose from the food, they were collected using a fine paintbrush and placed into a new vial containing a water moistened cotton ball. The larvae continued development at 21°C and upon eclosion and sclerotization, 50 individuals of each sex amongst all vials from each condition cohort and population were collected and stored in 70% ethanol for morphometric measurements.

#### Morphological measurements

Traits chosen for morphological measurement are based on previous research demonstrating their involvement in aggressive interactions. Primary among these is the foreleg (prothoracic leg) which has been shown to be involved with numerous aspects of aggressive behaviours such as thrusting, boxing/fencing and lunges (Dow and Schilcher 1975; Chen et al. 2002; Rohde et al. 2017; Dukas 2020). In addition to measurement of the forelegs, we also measured thorax length as a proxy for body size, as well as head width and wing length. Wing threats are used in aggressive displays (Chen et al. 2002), and there is evidence of a genetic correlation between displays and fighting in *Drosophila* (Baxter et al. 2023). However, there is no evidence that wings are used as weapons directly. As such, wing length would potentially differ in its response in contrast to the legs, which are directly used in physical combat. While the majority of aggressive interactions are among males (Jacobs 1960), females do sometimes display agonistic interactions with one another, often associated with the defense of a high-quality food resource (Ueda and Kidokoro 2002; Nilsen et al. 2004). Current evidence is not consistent with the outcome of female-female contests resulting in winner-loser hierarchies (Nilsen et al. 2004). While they share some of the same aggressive behaviours with males, their frequency of these differs substantially, and additionally will use “headbutting” (Nilsen et al. 2004; Zwarts et al. 2012), rarely seen in males. As such, head width was also included as a trait in our study.

Of the flies collected, 20 individuals of each sex of each cohort and treatment combination were dissected for imaging and subsequent measurement. Flies were dissected and images of the head, thorax, wing, and foreleg were taken with a Leica M125 stereoscope with a Leica DFC400 digital camera at magnifications of 50X or 63X, depending on the trait. Measurements of head width, thorax, wing length, wing width, femur, tibia, and first tarsal segment were conducted using ImageJ (1.53e) software (Rueden et al. 2017).

### Aggression Assays

To assess aggression, at generation 60 we removed 20 females from each treatment and replicate after the four days of territorial exposure. These females were allowed to lay eggs in two vials on two consecutive days, and density was controlled by culling excess eggs. After eclosion in each vial, 12 males were removed for the assay. Assays are described in Baxter and Dukas (2017) and summarised here. Two males of the same treatment were placed in arenas 3 cm in diameter with a patch of standard food 1.3 cm in diameter and a small ball of yeast and grapefruit juice. After the two males were added to the arenas, they were recorded for 15 minutes using a Logitech c920 camera. We ran 8 trials per lineage for a total of 32 trials for each of the 3 treatments. BORIS software (Friard & Gamba 2016) was used to score the footage with observers blind to fly treatment. Observers recorded wing threats, single male aggression (lunging or holding), and reciprocal male aggression (boxing or tussling).

### Statistical analyses

All analyses were done in R version 4.1.3. Response variables and the continuous predictor of thorax length were log_2_ transformed. The predictor variable of log_2_ (thorax length) was mean centred to aid model interpretation. Linear mixed models were fit using the glmmTMB version 1.1.4 package in R (Brooks et al. 2017). Both intercept and influence of thorax length were allowed to vary as random effects of replicate lineage nested within evolutionary treatment (thorax| Replicate). A similar model was also fit with starvation cohort as a predictor to determine if an interaction between the allometric coefficient and cohort existed to control for changes in allometry due to our starvation protocol. Plotting of observations identified several possible outliers, so models were run with and without outliers. Estimates were found to be similar, so outliers were included. Confidence intervals and *a priori*, custom contrasts used for inferences, were determined using emmeans version 1.8.0 (Lenth et al. 2018).

Data evaluating competitive fertilization success was modelled using a logistic generalized linear mixed model in glmmTMB with the counts of wild-type (“successes”) and crossveinless (“failures”) female offspring sired from a focal male as the response variable, and environmental treatment, male quality and their interaction as predictor variables. We also included random effects of experimental block and cage. We used emmeans to extract estimates and confidence intervals for treatment contrasts to our control treatment (NT). P-values from these estimates were adjusted using the Dunnett X method for two tests.

Data to evaluate changes in aggression were also modeled in glmmTMB, with evolutionary treatment and observer as fixed effects, while lineage nested within treatment, day of experiment and camera were modeled as independent random effects. Counts of lunges were modeled as Poisson with zero inflation. Threat duration is semi-continuous with zeros, and as such was modeled according to a Tweedie distribution (Tweedie power parameter estimated as ≍1.6). To confirm that the results (and the presence of many zeroes in the threat duration) were not unduly impacting model inferences (specifically treatment contrasts), we fit a similar model to the one above, but using a hurdle-Gamma, using the zero-inflated Gamma distribution in glmmTMB. This modeling strategy showed similar patterns of changes in aggression, of more modest magnitude. We used a log link for all these models. Visualization was done using ggplot2 (Wickham 2018).

## RESULTS

### Territorial restriction leads to increased fertilization success in high-quality males relative to non-territorial controls

To test how our territorial treatments influence variance in male reproductive success, we challenged both high- and low-quality focal males against marked tester males (crossveinless) in each territorial treatment. Consistent with our prediction, we observed an increase in competitive fertilization success in high quality males in the SCT treatment relative to the NT treatment (Figure 2; odds ratio (OR) of 4.25, 95% CIs 1.54-11.7, SE: 1.937, Z-ratio 3.17, p = 0.003). Consistent with the prediction there was a modest increase in siring success in the UCT treatment relative to the NT treatment (Figure 2; OR:3.09, CIs 1.01 −9.46, SE: 1.557, Z-ratio 2.24, p = 0.048). As expected, the differences in the environmental treatments had very modest influence on competitive siring success with low quality focal males (SCT/NT OR 0.98, CIs 0.34-2.79, SE:0.46, Z-ratio −0.05, p = 0.99; UCT/NT OR 0.24, CIs 0.06-0.89, SE:0.142, Z-ratio −2.41, p = 0.03). We also conducted an analysis of deviance (type II Wald χ^2^) for the interaction term between environmental treatment and male quality (χ^2^ = 8.78, df = 2, p = 0.012).

**Figure 2:**
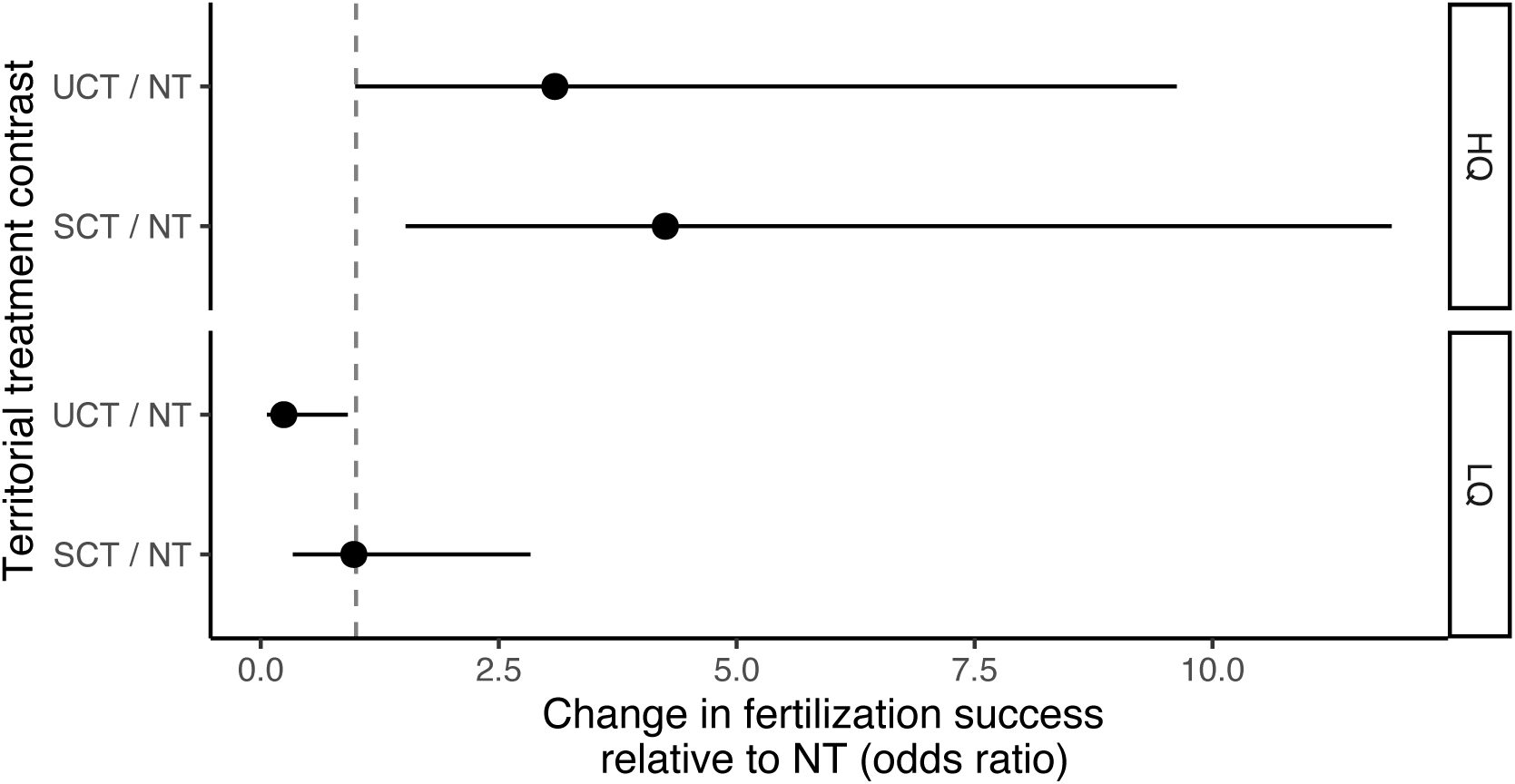
Odds ratio change between the territorial SCT and UCT treatments relative to the control NT treatment for both high- and low-quality male fertilization success against a crossveinless competitor. Model estimates plotted with 95% confidence intervals. HQ = high quality males, LQ = low quality males.

### Sexual size dimorphism (SSD) only mildly changed for some traits after 75 generations of experimental evolution

To test our prediction that territorial restriction would induce changes in sexual dimorphism due to intra-sexual competition, we measured Female-Male sexual dimorphism at generation 35 and 75 in all traits (Figure 3, S2, S3). We also modelled changes in SSD between condition cohorts as well as treatment (Figure 4). We observed no substantial changes in sexual dimorphism for any leg trait at either generation 35 (Figure 3A) or generation 75 (Figure 3B). At generation 75 we saw a change in SSD as a response to condition in the wing (Figure 4B). This effect appears to be due to a decrease in size of female wings. A similar effect was observed in femur length with a change in SSD when condition was accounted for (Figure 4B), that appeared to be due to a decrease in female femur size in UCT and SCT treatments in high condition but converging on similar trait values in low condition (Figure 4B). We also observed a change in SSD in tarsus when condition is accounted for (Figure 4B). Interestingly, this appears to be due to a lower condition response in tarsus in both sexes in UCT and SCT treatment, which is the opposite of the predicted trend (Figure 4B).

**Figure 3:**
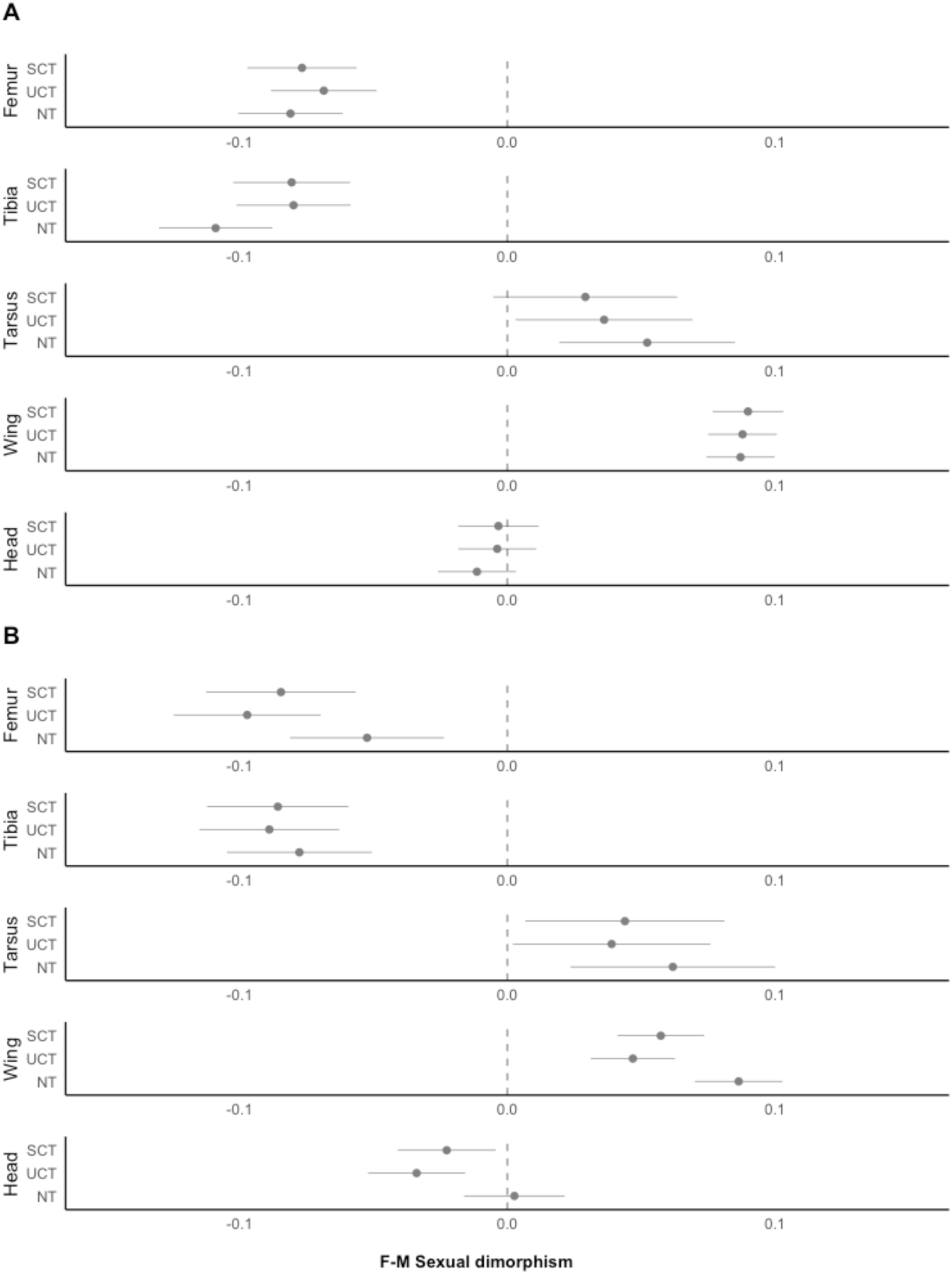
Trait and treatment specific sexual size dimorphism, for fully fed individuals, measured as difference between estimates of female – male trait sizes for (A) generation 35 and (B) generation 75. Response variables were log_2_ transformed. Model estimates plotted with 95% confidence intervals.

**Figure 4:**
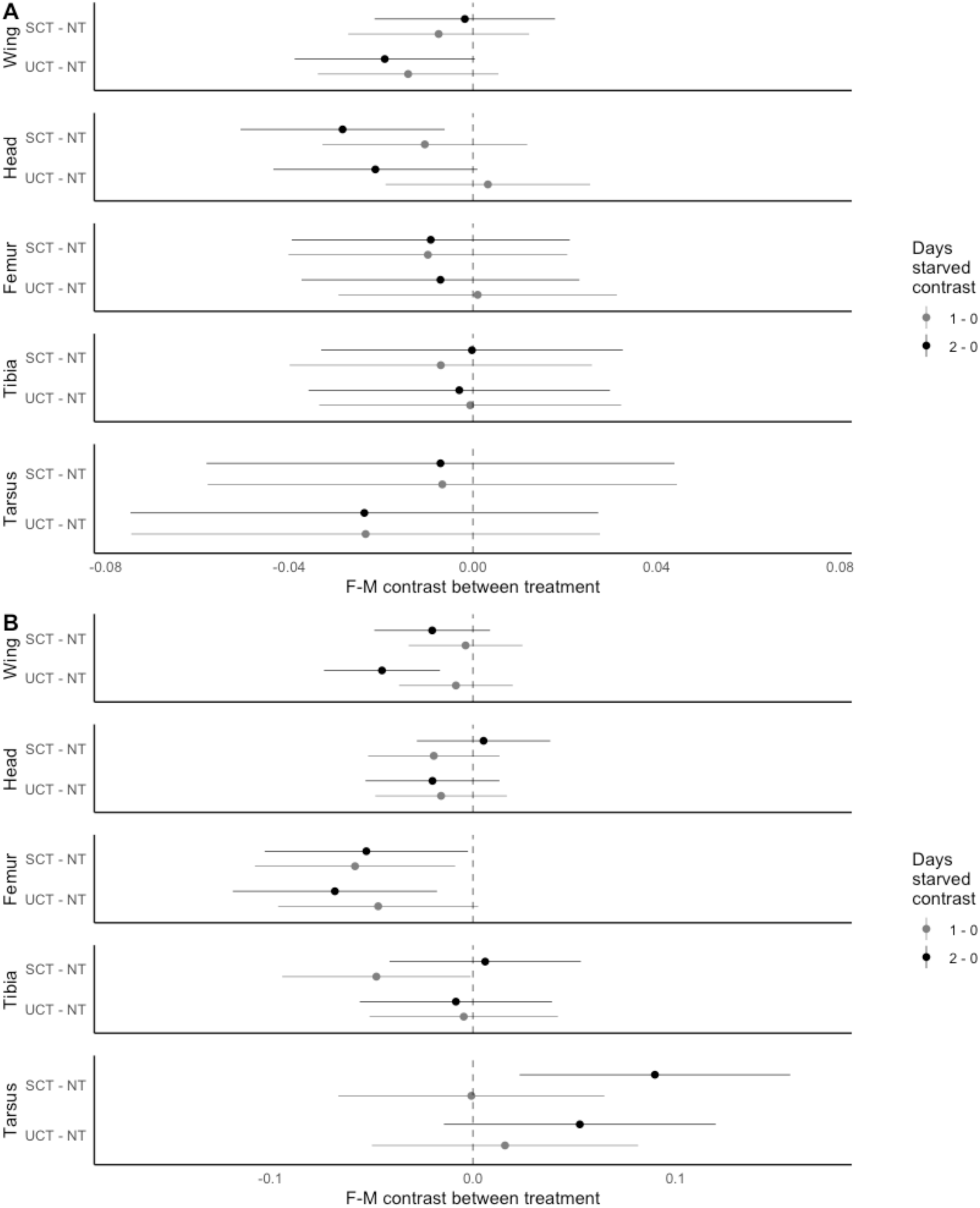
Trait specific changes among evolutionary treatments in sexual size dimorphism, relative to the NT treatment, measured as female trait size – male trait size for (A) generation 35 and (B) generation 75. Dimorphism was both contrasted between our non-restricted (NT) treatment and both territorial treatments (UCT and SCT) and food restriction condition treatment was contrasted with the non-restricted, high-quality treatment within each territorial treatment. Response variables were log_2_ transformed. Model estimates plotted with 95% confidence intervals.

### After 75 generations of experimental evolution, allometry changes were modest, and in the opposite direction of predictions

To test the prediction that the two territorially restricted treatments (UCT and SCT) would result in the evolution of increased allometric slopes (forelegs ∼ thorax) relative to NT, due to intra-sexual competition among males, we modeled the allometric slope for all measured traits at both generation 35 and generation 75 (Figure 5). At generation 35, the magnitude of changes in allometric relationships were modest for any trait relative to thorax size (Figure 5A, S4). Female head width in the SCT treatment had a lower allometric slope relative to NT, but with no concordant response observed in males (Figure 5A, S4). There was also some modest evidence for an interaction between sex, treatment, and head allometry in generation 35 (χ^2^=6.61, df=2, p=0.04). At generation 75 there were still no major evolutionary changes in the allometric slope for most traits (Figure 5B, S5). The allometric slope of femur seems to have decreased in females for both the SCT and UCT treatments relative to NT, but not in males (Figure 5B, S5), and there was a slightly significant sex by treatment by allometry interaction (χ^2^=7.34, df=2, p=0.03). The allometric slope of tarsus decreased in males of both UCT and SCT treatments, but not in females.

**Figure 5:**
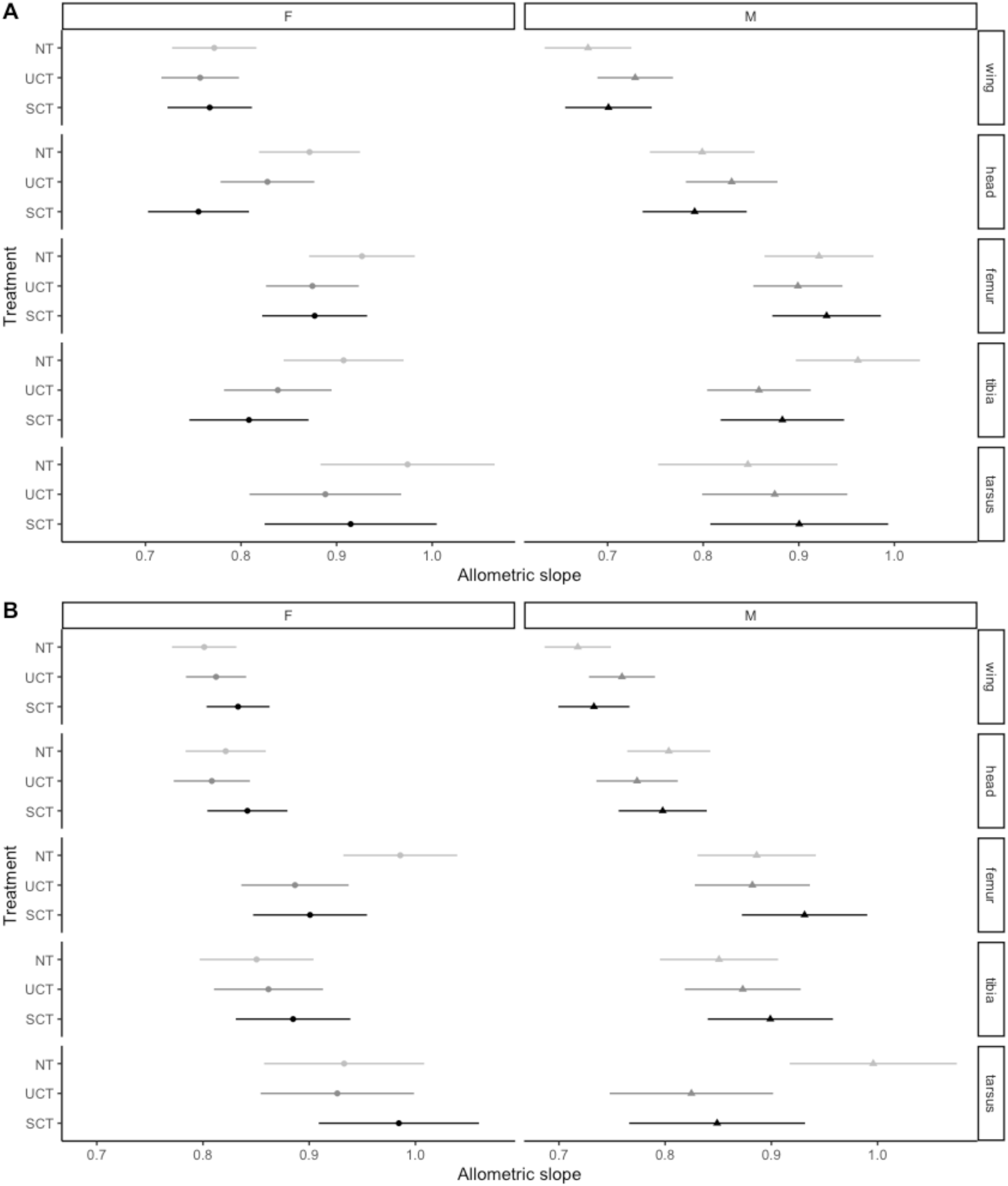
Allometric slopes for territorial treatments in each sex and each trait obtained from model estimated slopes of the response of trait size with log_2_ transformed thorax length as a predictor for (A) generation 35 and (B) generation 75. Bars represent 95% confidence intervals for model estimated slopes based on log_2_ transformed responses.

### After 75 generations, condition dependence in the tarsus of males decreased, with other traits showing inconsistent and modest responses

To test our prediction that an increase in condition dependence occurs concordantly with increases in sexual selection and trait exaggeration, we measured trait size responses to food deprivation in each treatment. At generation 35, all treatments and sexes showed the expected decrease in size with food deprivation (Figure 6A, S2). However, the slope of the response to manipulating condition shows very modest changes between treatments for most traits (Figure 6A). At generation 75, the overall reduction in size with deprivation is still observed (Figure 6B, S3). The response is slightly less compared to generation 35 (Figure 6A), but these experiments were done two years apart, performed by two separate individuals, so cannot be directly compared. The slope of the condition dependence appeared to increase for multiple traits in both sexes (a possible reduction of condition dependence) relative to NT (Figure 6). This increase was observed for wing size for both sexes, there was also an interaction between sex, treatment, and condition in the wing (χ^2^=8.30, df=2, p=0.016). There also appeared to be a minor decrease in condition dependence in the head in females. In the tarsus there was an interaction between sex, treatment, and condition dependence (χ^2^ =7.09, df=2, p=0.03). And both sexes seemed to be less condition dependent in the UCT treatment relative to the NT treatment (Figure 6). Males were also less condition dependent in the SCT treatment relative to NT (Figure 6).

**Figure 6:**
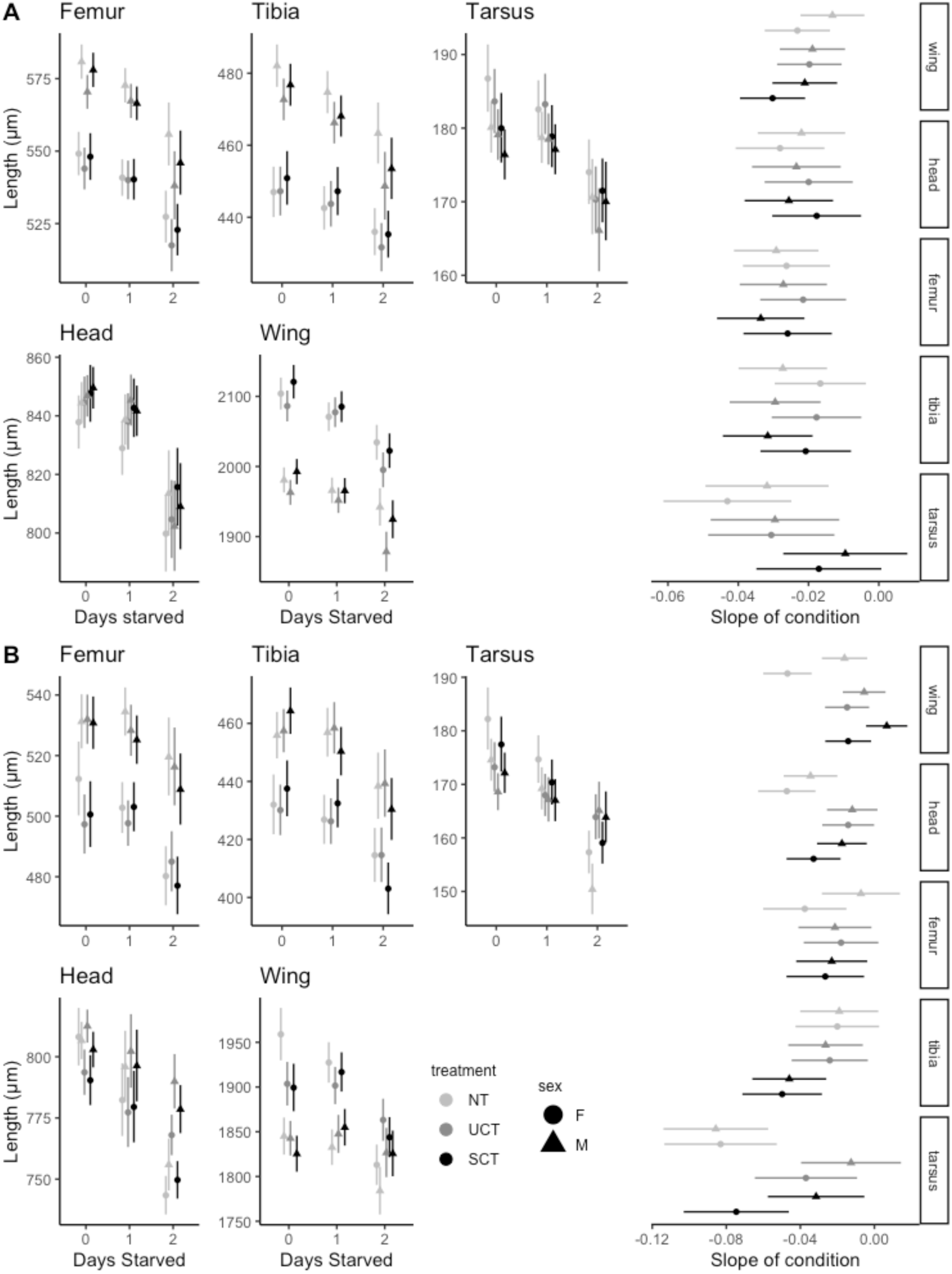
Condition response in trait size by treatment and sex and slope of response to condition by trait for (A) generation 35 and (B) generation 75. Trait size was log_2_ transformed for model fit, and back transformed for plotting. Estimates for trait size were run with condition as an ordinal predictor and used only for plotting, slope of condition was estimated with condition treatment as a continuous variable and was used for and size and condition inferences.

### There was modest increases in aggression in flies evolving in territorial treatments

At generation 60 there was an increase in the duration of threatening wing displays in males in the UCT and SCT treatment, (Figure 7A). There was also an increase in the number of lunges engaged by males of the UCT treatment relative to the NT treatment (Figure 7B). These responses both appear to be due to increases in the number of rare, highly aggressive males (Figure 7).

**Figure 7:**
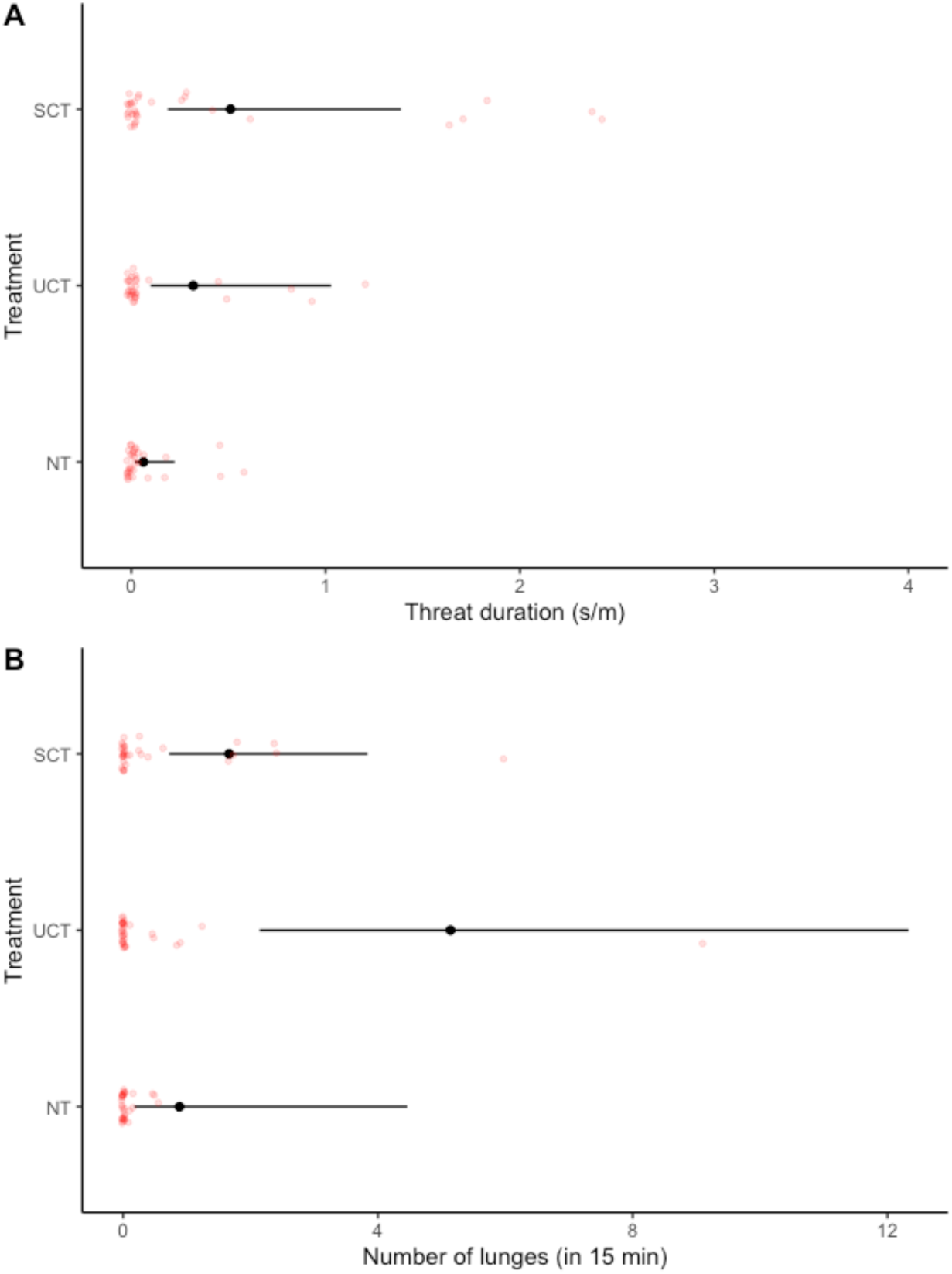
Estimated marginal means with 95% confidence intervals for aggression assays. (A) Estimated duration of threat displays for each treatment. In this panel, 2 values above 4 are not shown. (B) Model estimates for the number of lunges.

## DISCUSSION

According to the hypothesis laid out by Emlen (2008; 2014), the environmental precursors required for the evolution of weapons include: the possibility for males to defend a limiting resource required by females, intra-sexual competition favouring larger males or larger weapons, and these defendable and patchy resources are located in a way that facilitates one-on-one contests between males in which there is a winner and a loser. These three requirements should facilitate resource defense polygyny and differential reproductive success that is conducive to larger males who bear the largest weapons siring the bulk of the offspring. The experimental manipulations used in this study were developed to simulate these conditions in laboratory settings, and to partition some of the salient features regarding spatial constraints in access to the resources from overall defensibility and patchiness. The goal was to simulate conditions similar to what is observed in natural systems with resource defense polygyny, for instance, the guarding of tunnel entrances by dung beetle males (Emlen 1997), tree trunk fissures guarded by males and used as oviposition sites in antler flies (Dodson 1997), or the competition among males for overhanging rootlets with oviposition sites in stalk-eyed flies (Wilkinson and Dodson 1997). We demonstrate that these environmental conditions were sufficient to increase variance in male siring success with increasing territorial restriction, as predicted (Figure 2).

We predicted evolutionary increases in the allometric slopes in the legs of *D. melanogaster* males, generally used for intra-sexual combat (Dow and Schilcher 1975; Chen et al. 2002; Rohde et al. 2017; Dukas 2020), in our UCT and SCT treatments, due to increased opportunity for territorial defence and increased mating success for males who defend a territory. While we did observe “statistically significant” evolutionary changes in morphology, broadly speaking, the results from our experiment were not consistent with these predictions. We did not observe consistent evolutionary increases in male biased dimorphism in the legs, allometric coefficients did not evolve substantially, and decreased for some traits, in all treatments compared to our non-territorial (NT) control treatment (Figure 5). Recent work has argued that ‘pure’ weapons (used solely for combat), but not for threat signaling, may not be under selection for increases in allometric slope, whereas “threat signals” would evolve such a response (McCullough and O’Brien 2022). In addition to its locomotory role, the legs of *D. melanogaster* are used in physical combat, but not to our knowledge as a threat signal. However, this weapon-signal continuum suggests aggressive signalling traits, such as wings in *D. melanogaster*, would be under selection for increased allometric slope, which we also do not observe (Figure 5; Chen et al. 2002; McCullough and O’Brien 2022). We highlight the importance of working from pre-established hypotheses and predictions in experimental evolution studies, as these experiments are conducive to evolutionary changes (as we observed), potentially unrelated to the purpose of the experiment.

Exaggerated male traits (such as weapons) often display sensitivity to condition proportionally greater than non-exaggerated traits, an increase in condition dependence (Rowe and Houle 1996; Bonduriansky and Day 2003). This increased condition dependence in exaggerated traits has been hypothesised to occur as a signal of overall quality. Larger weapons are a possible indicator that a male has high quality alleles that can produce a large trait (Zahavi 1975; Andersson 1994; Kirkpatrick 1996). This would of course mean that exaggerated sexual traits are under constant directional selection, and the genetic variation would rapidly deplete, known as the lek paradox. However, the genic capture model proposes that as trait exaggeration occurs, many loci become associated with trait expression, maintaining genetic variation for the trait (Rowe and Houle 1996). This would imply that trait expression may be representative of both genetic quality and resource acquisition due to the contribution of much of the genome. Because of this consistent pattern of condition dependence in sexually selected weapons and ornaments, we would also expect increases in male foreleg condition dependence in our territorial restriction. Contrary to these predictions, we found increased condition dependence in female traits and decreases in condition dependence in males (Figure 6, S5). This reduced condition dependence was observed in the wing, head, and tarsus with no change in the femur or tibia which were the traits predicted to respond to territorial selection. Although this runs counter to our prediction, even if we did see trait exaggeration it is not known at what stage we would expect condition dependence to evolve. Under a good genes model where where variation for the indicator trait is oligogenic, condition dependence may evolve later in the evolution of trait exaggeration once the trait has already begun to increase in size and potentially after a depletion of the segregating variation can be used as an indicator of quality. It may also be the case that trait exaggeration occurs most often for traits that already show heightened condition dependence because they already act as reliable indicators before exaggeration, as suggested by Johnstone et al. (2009). In this case, we would not expect to observe changes in condition dependence with increased sexual selection or SSD for the forelegs. The genic capture model suggests that many alleles will be responsible for the trait, and that many alleles associated will be co-opted by these trait loci (Rowe and Houle 1996). Because the forelegs have additional locomotory functions, and their composite nature has been historically shaped by natural selection, it may be more difficult to co-opt condition genes into trait expression, in comparison to a novel trait like a beetle horn. Despite the additional locomotory fuctions however, a number of insects do display exaggerated legs (Singh 1977; Zeh et al. 1992).

If Emlen’s (2014) hypothesis for the three key conditions for weapon evolution, and the evolution of condition dependence to maintain sexually selected traits is true, there are experimental reasons we may have failed to detect morphological change that should be considered. Primarily amongst them is the density of individuals in our experiment. Although Hoffmann and Cacoyianni (1990) demonstrated resource defense polygyny in *D. melanogaster* at a similar resource patch diameter used in this experiment, they also found that at higher densities, defense was abandoned and flies reverted to scramble competition. Our findings may suggest that for trait exaggeration to initiate, populations may require generally low density to avoid interactions where resource defense is not a viable option due to pressures from multiple competitors at once. If we limited the census size, however, it would necessarily reduce genetic variation in our experiment and possibly diminish the likelihood of selecting on alleles of interest. The other possible experimental impact is also due to uncontrolled density creating possible limits on food for larvae. Although we provided both high-quality food as well as a low-quality food resource for adults, larvae in the high-quality food appeared to be growing under high density conditions in the food. This high larval density may have created additional countervaling selection pressure on size due to larval competition. Although we did not test fecundity, the observation that density was high may suggest that fecundity selection and larval viability selection predominated, rather than sexual selection on males. With very high-quality food provided and a high adult density, there may have been a strong enough selection pressure on females to lay as many eggs as early as possible to ‘beat the rush’. This could counter impacts of sexual selection on males in the treatments. In a previous study using similar spatial-environment treatments to assess selective dynamics of the purging of deleterious alleles, it was observed that the impact of the spatial-environment varied among deleterious mutations influencing a variety of traits (Wilson et al. 2021). As such, the three environmental manipulations we used to influence opportunity for sexual selection, may have also influenced other components of natural selection to varying degrees.

A large portion of the literature has been dedicated to the idea that weapons or traits used for resource defense develop positive allometries (allometric coefficient >1; Kodric-Brown et al. 2006; Eberhard et al. 2018). This has been shown in a number of species including beetle species (Kawano 1995), cervids (Lemaître et al. 2014), stalk-eyed flies (Baker and Wilkinson 2001), and many others (Voje 2016). While weapons are often associated with positive allometries, it is by no means always the case (Voje 2016). In our study, the allometry of leg traits are generally the greatest in magnitude (although with slopes still less than 1), in comparison to other traits. Femur length, while phenotypically plastic, scales approximately isometrically with overall size in response to changes in density or food availability (Shingleton et al. 2009; Pesevski 2021). Although we saw a general trend towards smaller size and lower allometric coefficients in the legs, confidence intervals often overlapped substantially across treatments and both sexes, with generally small magnitudes across evolutionary treatments (slope changes). This may simply reflect the constraints to sexually dimorphic evolution due to genetic correlation (r_MF_) between the sexes (Lande 1980). Previous studies applying strong, sexually discordant, mass artificial selection on males and female body size, found it took more than 100 generations to observe sex-specific responses, while sex-concordant responses were much faster (Stewart and Rice 2018; Audet et al. 2024). This suggests that 75 generations of what is likely modest selective pressure (in terms of changes to sexual selection *per se*, as a result of the spatial environments employed), may have been insufficient number of generations for a sex-specific response (Stewart and Rice 2018). In plants it has been shown that the breakdown of r_MF_ can result in rapid changes in SD (Delph et al. 2011), but this reduced constraint was imposed by strong family-based selection. In general, homologous traits between the sexes have a very high genetic correlation, providing some constraints on the potential to evolve dimorphically (Poissant et al. 2010). The legs in *D. melanogaster* although used in intra-sexual competition even under all the predicted circumstances proposed by Emlen, are still constrained by the shared genome, the biomechanical function of legs in locomotion, and the time required to adapt morphological changes with a shared genome with experimental evolution may be significantly longer than what is experimentally feasible. This is of course opposed by the fact that there was at least a small change in SSD in both head and wing, but not the focal trait of leg. r_MF_ is trait specific and response to sexually discordant selection occurs in a trait specific manner, which has been previously suggested (Poissant et al. 2010; Audet et al. 2024).

The one trait that we observed to respond in a consistently sex specific way was the tarsus of males (Figure 5B, 5B). It appears as though after 75 generations of evolution, male tarsi have become more hypo-allometric and less condition dependent. This may be due to the sex-limited structure on the tarsi of males, the sex comb. The presence of sex combs *in D. melanogaster* is important for mating success (Ng and Kopp 2008), and the length of the tarsus is directly related to sex comb number (Combs 1937). For this reason, the presence and necessity of sex combs may be creating a physical barrier to decreased size evolution and condition dependent response in males.

The results here, although not aligned with either Emlen’s hypotheses for weapon evolution or the condition dependence hypothesis, are consistent with other experimental evolution studies. These experiments show results inconsistent with previous hypotheses for responses to experimental evolution when sexual selection dynamics are altered via manipulating sex ratio (Bath et al. 2021, 2023; Edmunds et al. 2021; Sepil et al. 2022). By manipulating census sex ratio, Bath et al. (2021, 2023) and Sepil et al. (2022) attempted to modulate the intensity of sexual selection in a long-term experimental population to explore the potential consequences of heightened sexual selection. Consistent with our observations (Figure 7), these experiments found modest increases in male aggression with increased sexual competition and a more substantial increase in female post mating aggression (Bath et al. 2021). They also observed no change in male condition dependence but increased female condition dependence. Despite a very different experimental approach, their results, similar to ours, show that in the context of heightened sexual selection, male condition dependence does not increase, while female condition dependence does (Bath et al. 2023). In male biased sex ratio populations (a proxy for increased sexual competition) it appeared that males decreased investment in pre-copulatory investment, but also did not increase mating duration which would be suggestive of post-copulatory investments (Sepil et al. 2022). These results combined with ours may suggest that the condition dependence model of sexually dimorphic evolution may rely on assumptions of a relatively simple genetic architecture, and that quantitative traits may require very strong and consistent directional selection to show a response. This has been suggested previously, and even modelled. Johnstone et al. (2009) suggest that strong sexual selection for an increase in trait size does not require the evolution of condition dependence, and if it does, this may be a brief increase in condition dependence that reduces over time. Our results as well as Bath et al. (2023), suggest that the evolution of condition dependence is not a necessary mechanism to maintain genetic variation for traits under persistent sexual selection.

## Supporting information

All supplemental figures and tables

## Author contributions

Study Conceptualization and funding: ID

Study Design: ID, AW

Artificial Selection and rearing: AW, TA

Male Mating Success Experiment: AW

Aggression experiment: RD

Image Analysis: AW, TA

Analysis: ID, AW, TA

Manuscript drafting: TA, AW, ID

Manuscript editing: TA, AW, ID, RD

Manuscript revisions: ID, TA, RD, AW

## Conflict of Interest statement

The authors declare no conflict of interests.

Upon acceptance all data and scripts will be made available via github as well as either Figshare or DRYAD.

## Acknowledgements and Funding

We thank Francesco Ruso for dissecting. This work was funded by NSERC Discovery and Discovery Accelerator grants to ID and RD and Ontario Graduate Scholarships to TA and AW.

## REFERENCES

1. Andersson, M. 1994. Sexual Selection. Princeton University Press.

2. Audet, T., J. Krol, K. Pelletier, A. D. Stewart, and I. Dworkin. 2024. Sexually discordant selection is associated with trait-specific morphological changes and a complex genomic response. Evolution qpae071.

3. Baker, R. H., and G. S. Wilkinson. 2001. Phylogenetic Analysis of Sexual Dimorphism and Eye-Span Allometry in Stalk-Eyed Flies (diopsidae). Evolution 55:1373–1385.

4. Bateman, A. J. 1948. Intra-sexual selection in Drosophila. Heredity 2:349–368.

5. Bath, E., D. Edmunds, J. Norman, C. Atkins, L. Harper, W. G. Rostant, T. Chapman, S. Wigby, and J. C. Perry. 2021. Sex ratio and the evolution of aggression in fruit flies. Proceedings of the Royal Society B: Biological Sciences 288:20203053. Royal Society.

6. Bath, E., W. Rostant, H. J. Ostridge, S. Smith, J. S. Mason, T. Rafaluk-Mohr, J. E. Mank, T. Chapman, and J. C. Perry. 2023. Sexual selection and the evolution of condition-dependence: an experimental test at two resource levels. Evolution 77:776–788.

7. Baxter, C., and R. Dukas. 2017. Life history of aggression: effects of age and sexual experience on male aggression towards males and females - ScienceDirect.

8. Baxter, C. M., I. Shams, I. Dworkin, and R. Dukas. 2023. Genetic correlation between aggressive signals and fighting. Biology Letters 19:20220616. Royal Society.

9. Baxter, C., J. Mentlik, I. Shams, and R. Dukas. 2018. Mating success in fruit flies: courtship interference versus female choice. Animal Behaviour 138:101–108.

10. Boake, C. R. B., M. P. DeAngelis, and D. K. Andreadis. 1997. Is sexual selection and species recognition a continuum? Mating behavior of the stalk-eyed fly Drosophila heteroneura. Proceedings of the National Academy of Sciences 94:12442–12445. Proceedings of the National Academy of Sciences.

11. Boisseau, R. P., M. M. Ero, S. Makai, L. J. G. Bonneau, and D. J. Emlen. 2020. Sexual dimorphism divergence between sister species is associated with a switch in habitat use and mating system in thorny devil stick insects. Behavioural Processes 181:104263.

12. Bonduriansky, R. 2007. The Evolution of Condition Dependent Sexual Dimorphism. 11.

13. Bonduriansky, R., and T. Day. 2003. The Evolution of Static Allometry in Sexually Selected Traits. Evolution 57:2450–2458.

14. Brooks, M., E., K. Kristensen, K. Benthem, J.,van, A. Magnusson, C. Berg W., A. Nielsen, H. Skaug J., M. Mächler, and B. Bolker M. 2017. glmmTMB Balances Speed and Flexibility Among Packages for Zero-inflated Generalized Linear Mixed Modeling. The R Journal 9:378.

15. Ceballos, C. P., and N. Valenzuela. 2011. The Role of Sex-specific Plasticity in Shaping Sexual Dimorphism in a Long-lived Vertebrate, the Snapping Turtle Chelydra serpentina. Evol Biol 38:163.

16. Chen, S., A. Y. Lee, N. M. Bowens, R. Huber, and E. A. Kravitz. 2002. Fighting fruit flies: A model system for the study of aggression. Proceedings of the National Academy of Sciences 99:5664–5668. Proceedings of the National Academy of Sciences.

17. Combs, J. D. 1937. Genetic and Environmental Factors Affecting the Development of the Sex-Combs of Drosophila Melanogaster. Genetics 22:427–433.

18. Darwin, C. 1871. The Descent of Man, and Selection in Relation to Sex. D. Appleton.

19. David, P., T. Bjorksten, K. Fowler, and A. Pomiankowski. 2000. Condition-dependent signalling of genetic variation in stalk-eyed flies. Nature 406:186–188. Nature Publishing Group.

20. Delph, L. F., J. C. Steven, I. A. Anderson, C. R. Herlihy, and E. D. Brodie III. 2011. Elimination of a Genetic Correlation Between the Sexes Via Artificial Correlational Selection. Evolution 65:2872–2880.

21. Dodson, G. N. 1997. Resource Defense Mating System in Antlered Flies, Phytalmia spp. (Diptera: Tephritidae). Annals of the Entomological Society of America 90:496–504.

22. Dow, M. A., and F. V. Schilcher. 1975. Aggression and mating success in Drosophila melanogaster. Nature 254:511–512. Nature Publishing Group.

23. Dukas, R. 2020. Natural history of social and sexual behavior in fruit flies. Sci Rep 10:21932. Nature Publishing Group.

24. Dweck, H. K. M., S. A. M. Ebrahim, S. Kromann, D. Bown, Y. Hillbur, S. Sachse, B. S. Hansson, and M. C. Stensmyr. 2013. Olfactory Preference for Egg Laying on Citrus Substrates in Drosophila. Current Biology 23:2472–2480.

25. Eberhard, W. G., R. L. Rodríguez, B. A. Huber, B. Speck, H. Miller, B. A. Buzatto, and G. Machado. 2018. Sexual Selection and Static Allometry: The Importance of Function. The Quarterly Review of Biology 93:207–250.

26. Edmunds, D., S. Wigby, and J. C. Perry. 2021. A resource-poor developmental diet reduces adult aggression in male Drosophila melanogaster. Behav Ecol Sociobiol 75:110.

27. Emberts, Z., W. S. Hwang, and J. J. Wiens. 2021. Weapon performance drives weapon evolution. Proceedings of the Royal Society B: Biological Sciences 288:20202898. Royal Society.

28. Emlen, D. J. 1997. Alternative reproductive tactics and male-dimorphism in the horned beetle Onthophagus acuminatus (Coleoptera: Scarabaeidae). Behavioral Ecology and Sociobiology 41:335–341.

29. Emlen, D. J. 2014. Reproductive contests and the evolution of extreme weaponry. Oxford University Press.

30. Emlen, D. J. 2008. The Evolution of Animal Weapons. Annual Review of Ecology, Evolution, and Systematics 39:387–413.

31. Emlen, D. J., and T. K. Philips. 2006. Phylogenetic Evidence for an Association Between Tunneling Behavior and the Evolution of Horns in Dung Beetles (Coleoptera: Scarabaeidae: Scarabaeinae). cole 60:47–56. The Coleopterists Society.

32. Emlen, S. T., and L. W. Oring. 1977. Ecology, Sexual Selection, and the Evolution of Mating Systems. Science 197:215–223. American Association for the Advancement of Science.

33. Fairbairn, D. J. 2005. Allometry for Sexual Size Dimorphism: Testing Two Hypotheses for Rensch’s Rule in the Water Strider Aquarius remigis. 16.

34. Guo, X., and R. Dukas. 2020. The cost of aggression in an animal without weapons. Ethology 126:24–31.

35. Harshman, L. G., and A. A. Hoffmann. 2000. Laboratory selection experiments using Drosophila: what do they really tell us? Trends in Ecology & Evolution 15:32–36. Elsevier.

36. Hoffmann, A. A., and Z. Cacoyianni. 1989. Selection for territoriality in *Drosophila melanogaster*: correlated responses in mating success and other fitness components. Animal Behaviour 38:23–34.

37. Hoffmann, A. A., and Z. Cacoyianni. 1990. Territoriality in Drosophila melanogaster as a conditional strategy. Animal Behaviour 40:526–537.

38. Honěk, A. 1993. Intraspecific Variation in Body Size and Fecundity in Insects: A General Relationship. Oikos 66:483–492. [Nordic Society Oikos, Wiley].

39. Hurd, P. L. 2006. Resource holding potential, subjective resource value, and game theoretical models of aggressiveness signalling. Journal of Theoretical Biology 241:639–648.

40. Jacobs, M. E. 1960. Influence of Light on Mating of Drosophila Melanogaster. Ecology 41:182–188. Ecological Society of America.

41. Jensen, K., C. McClure, N. K. Priest, and J. Hunt. 2015. Sex-specific effects of protein and carbohydrate intake on reproduction but not lifespan in Drosophila melanogaster. Aging Cell 14:605–615.

42. Johns, A., H. Gotoh, E. L. McCullough, D. J. Emlen, and L. C. Lavine. 2014. Heightened Condition-Dependent Growth of Sexually Selected Weapons in the Rhinoceros Beetle, Trypoxylus dichotomus (Coleoptera: Scarabaeidae). Integrative and Comparative Biology 54:614–621.

43. Johnstone, R. A., S. A. Rands, and M. R. Evans. 2009. Sexual selection and condition-dependence. Journal of Evolutionary Biology 22:2387–2394.

44. Kawano, K. 1995. Horn and Wing Allometry and Male Dimorphism in Giant Rhinoceros Beetles (Coleoptera: Scarabaeidae) of Tropical Asia and America. Annals of the Entomological Society of America 88:92–99.

45. Kirkpatrick, M. 1996. Good Genes and Direct Selection in the Evolution of Mating Preferences. Evolution 50:2125–2140.

46. Kodric-Brown, A., R. M. Sibly, and J. H. Brown. 2006. The allometry of ornaments and weapons. Proceedings of the National Academy of Sciences 103:8733–8738.

47. Kokko, H., R. Brooks, J. M. McNamara, and A. I. Houston. 2002. The sexual selection continuum. Proc. R. Soc. Lond. B 269:1331–1340.

48. Kudo, A., S. Shigenobu, K. Kadota, M. Nozawa, T. F. Shibata, Y. Ishikawa, and T. Matsuo. 2017. Comparative analysis of the brain transcriptome in a hyper-aggressive fruit fly, Drosophila prolongata. Insect Biochemistry and Molecular Biology 82:11–20.

49. Lande, R. 1980. Sexual Dimorphism, Sexual Selection, and Adaptation in Polygenic Characters. Evolution 34:292–305. [Society for the Study of Evolution, Wiley].

50. Lee, K. P., S. J. Simpson, F. J. Clissold, R. Brooks, J. W. O. Ballard, P. W. Taylor, N. Soran, and D. Raubenheimer. 2008. Lifespan and reproduction in Drosophila: New insights from nutritional geometry. Proceedings of the National Academy of Sciences 105:2498–2503. Proceedings of the National Academy of Sciences.

51. Lemaître, J. F., C. Vanpé, F. Plard, and J. M. Gaillard. 2014. The allometry between secondary sexual traits and body size is nonlinear among cervids. Biology Letters 10:20130869. Royal Society.

52. Lenth, R., H. Singmann, J. Love, and P. Buerkner. 2018. emmeans: Estimated marginal means, aka Least-Squares Means.

53. Lindenfors, P., C. L. Nunn, and R. A. Barton. 2007. Primate brain architecture and selection in relation to sex. BMC Biology 5:20.

54. Maklakov, A. A., S. J. Simpson, F. Zajitschek, M. D. Hall, J. Dessmann, F. Clissold, D. Raubenheimer, R. Bonduriansky, and R. C. Brooks. 2008. Sex-Specific Fitness Effects of Nutrient Intake on Reproduction and Lifespan. Current Biology 18:1062–1066.

55. McCullough, E. L., C. W. Miller, and D. J. Emlen. 2016. Why Sexually Selected Weapons Are Not Ornaments. Trends in Ecology & Evolution 31:742–751.

56. McCullough, E. L., and D. M. O’Brien. 2022. Variation in allometry along the weapon-signal continuum. Evol Ecol 36:591–604.

57. Moczek, A. P., and D. J. Emlen. 2000. Male horn dimorphism in the scarab beetle, Onthophagus taurus: do alternative reproductive tactics favour alternative phenotypes? Animal Behaviour 59:459–466.

58. Nandy, B., V. Gupta, S. Sen, N. Udaykumar, M. A. Samant, S. Z. Ali, and N. G. Prasad. 2013. Evolution of mate-harm, longevity and behaviour in male fruit flies subjected to different levels of interlocus conflict. BMC Evolutionary Biology 13:212.

59. Ng, C. S., and A. Kopp. 2008. Sex Combs are Important for Male Mating Success in Drosophila melanogaster. Behav Genet 38:195–201.

60. Nilsen, S. P., Y.-B. Chan, R. Huber, and E. A. Kravitz. 2004. Gender-selective patterns of aggressive behavior in Drosophila melanogaster. Proceedings of the National Academy of Sciences 101:12342–12347. Proceedings of the National Academy of Sciences.

61. Palaoro, A. V., and P. E. C. Peixoto. 2022. The hidden links between animal weapons, fighting style, and their effect on contest success: a meta-analysis. Biological Reviews 97:1948–1966.

62. Partridge, L., A. Ewing, and A. Chandler. 1987. Male size and mating success in *Drosophila melanogaster*: the roles of male and female behaviour. Animal Behaviour 35:555–562.

63. Perdigón Ferreira, J., P. T. Rohner, and S. Lüpold. 2023. Strongly sexually dimorphic forelegs are not more condition-dependent than less dimorphic traits in Drosophila prolongata. Evol Ecol, doi: 10.1007/s10682-022-10226-0.

64. Pesevski, M. 2021. Influence of environmental variation on sexual dimorphism in Drosophila morphology among adaptively diverged populations and in an inter-specific comparative context. (Doctoral dissertation).

65. Poissant, J., A. J. Wilson, and D. W. Coltman. 2010. Sex-Specific Genetic Variance and the Evolution of Sexual Dimorphism: A Systematic Review of Cross-Sex Genetic Correlations. Evolution 64:97–107.

66. Reddiex, A. J., T. P. Gosden, R. Bonduriansky, and S. F. Chenoweth. 2013. Sex-Specific Fitness Consequences of Nutrient Intake and the Evolvability of Diet Preferences. The American Naturalist 182:91–102. The University of Chicago Press.

67. Reeve, J. P., and D. J. Fairbairn. 1999. Change in sexual size dimorphism as a correlated response to selection on fecundity. Heredity 83:697–706. Nature Publishing Group.

68. Rohde, P. D., B. Gaertner, K. Ward, P. Sørensen, and T. F. C. Mackay. 2017. Genomic Analysis of Genotype-by-Social Environment Interaction for Drosophila melanogaster Aggressive Behavior. Genetics 206:1969–1984.

69. Rowe, L., and D. Houle. 1996. The lek paradox and the capture of genetic variance by condition dependent traits. Proc. R. Soc. Lond. B 263:1415–1421.

70. Rueden, C. T., J. Schindelin, M. C. Hiner, B. E. DeZonia, A. E. Walter, E. T. Arena, and K. W. Eliceiri. 2017. ImageJ2: ImageJ for the next generation of scientific image data. BMC Bioinformatics 18:529.

71. Scott, A. M., I. Dworkin, and R. Dukas. 2022. Evolution of sociability by artificial selection*. Evolution 76:541–553.

72. Sepil, I., J. C. Perry, A. Dore, T. Chapman, and S. Wigby. 2022. Experimental evolution under varying sex ratio and nutrient availability modulates male mating success in Drosophila melanogaster. Biology Letters 18:20210652. Royal Society.

73. Shingleton, A. W., C. M. Estep, M. V. Driscoll, and I. Dworkin. 2009. Many ways to be small: different environmental regulators of size generate distinct scaling relationships in Drosophila melanogaster. Proceedings of the Royal Society B: Biological Sciences 276:2625–2633. Royal Society.

74. Simmons, L. W., and J. L. Tomkins. 1996. Sexual selection and the allometry of earwig forceps. Evol Ecol 10:97–104.

75. Singh, B. 1977. Two new and two unrecorded species of the genus Drosophila Fallen (Diptera: Drosophilidae) from Shillong, Meghalaya, India.

76. Spieth, H. T. 1974. Courtship Behavior in Drosophila. Annual Review of Entomology 19:385–405. Annual Reviews.

77. Spieth, H. T. 1981. Drosophila heteroneura and Drosophila silvestris: Head Shapes, Behavior and Evolution. Evolution 35:921–930. [Society for the Study of Evolution, Wiley].

78. Stewart, A. D., and W. R. Rice. 2018. Arrest of sex-specific adaptation during the evolution of sexual dimorphism in Drosophila. Nat Ecol Evol 2:1507–1513.

79. Stillwell, R. C., I. Dworkin, A. W. Shingleton, and W. A. Frankino. 2011. Experimental Manipulation of Body Size to Estimate Morphological Scaling Relationships in Drosophila. J Vis Exp 3162.

80. Summers, T. C., and T. J. Ord. 2022. Female preference for super-sized male ornaments and its implications for the evolution of ornament allometry. Evol Ecol 36:701–716.

81. Tanaka, T., and T. Yamazaki. 1990. Fitness and its components in Drosophila melanogaster. 遺伝学雑誌 65:417–426.

82. Tatar, M. 2011. The plate half-full: Status of research on the mechanisms of dietary restriction in Drosophila melanogaster. Experimental Gerontology 46:363–368.

83. Taylor, P. D., and G. C. Williams. 1982. The lek paradox is not resolved. Theoretical Population Biology 22:392–409.

84. Toyoshima, N., and T. Matsuo. 2023. Fight outcome influences male mating success in Drosophila prolongata. J Ethol, doi: 10.1007/s10164-023-00778-1.

85. Ueda, A., and Y. Kidokoro. 2002. Aggressive behaviours of female Drosophila melanogaster are influenced by their social experience and food resources. Physiological Entomology 27:21–28.

86. Voje, K. L. 2016. Scaling of Morphological Characters across Trait Type, Sex, and Environment: A Meta-analysis of Static Allometries. The American Naturalist 187:89–98.

87. Wickham, H. 2018. ggplot2: Elegant Graphics for Data Analysis. Springer-Verlag New York, New York.

88. Wilkinson, G. S., and G. N. Dodson. 1997. Function and evolution of antlers and eye stalks in flies. Pp. 310–328 in J. C. Choe and B. J. Crespi, eds. The Evolution of Mating Systems in Insects and Arachnids. Cambridge University Press.

89. Wilson, A. E., A. Siddiqui, and I. Dworkin. 2021. Spatial heterogeneity in resources alters selective dynamics in Drosophila melanogaster. Evolution 75:1792–1804.

90. Zahavi, A. 1975. Mate selection—A selection for a handicap. Journal of Theoretical Biology 53:205–214.

91. Zeh, D. W., J. A. Zeh, and G. Tavakilian. 1992. Sexual Selection and Sexual Dimorphism in the Harlequin Beetle Acrocinus longimanus. Biotropica 24:86–96. [Association for Tropical Biology and Conservation, Wiley].

92. Zwarts, L., M. Versteven, and P. Callaerts. 2012. Genetics and neurobiology of aggression in Drosophila. Fly 6:35–48. Taylor & Francis.

