## Supplementary material for "The role of resource defensibility in facilitating sexually-selected weapon evolution: An experimental evolution test": All supplemental figures and tables

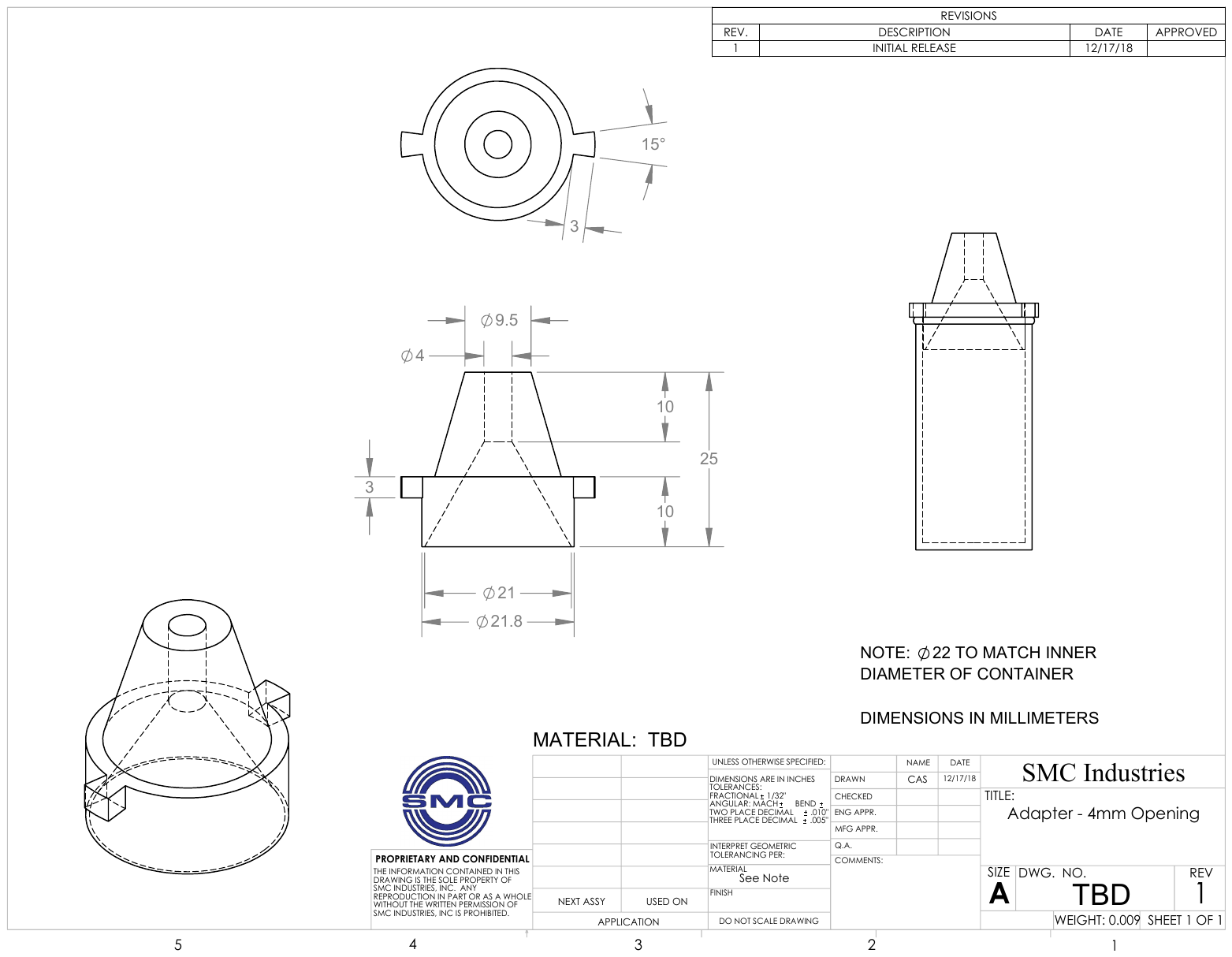

Figure S1: Schematic drawing of entrance restriction caps that were used for the SCT treatment. Caps were built of hard plastic. All dimension displayed in millimeters.

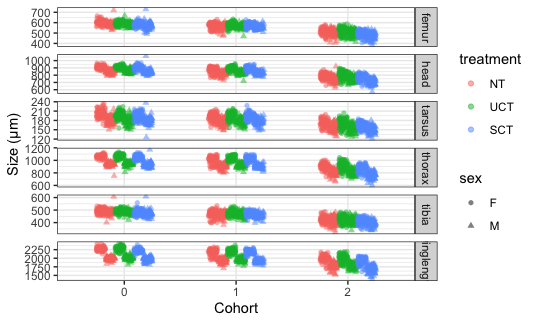

Figure S2: Generation 35 unmodelled size measurements for all measured traits as with sexes and evolutionary treatment separate.

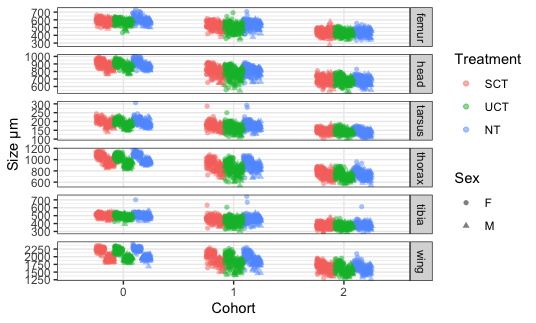

Figure S3: Generation 75 unmodelled size measurements for all measured traits as with sexes and evolutionary treatment separate.

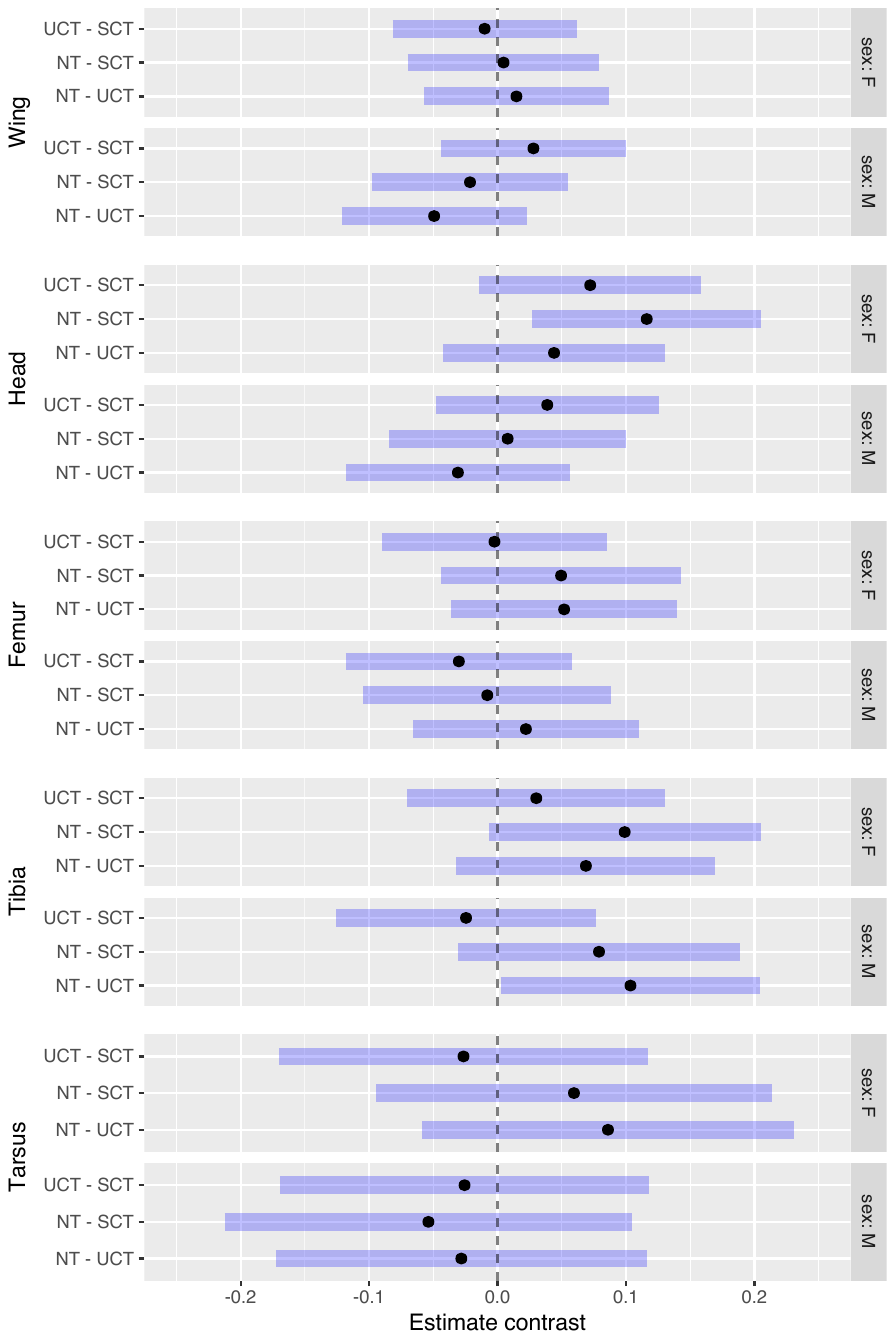

Figure S4: Generation 35 Allometry contrasts, and their 95% confidence intervals from model estimates. Measurements were log_2_ transformed before modelling and contrasts were computed with emmeans.

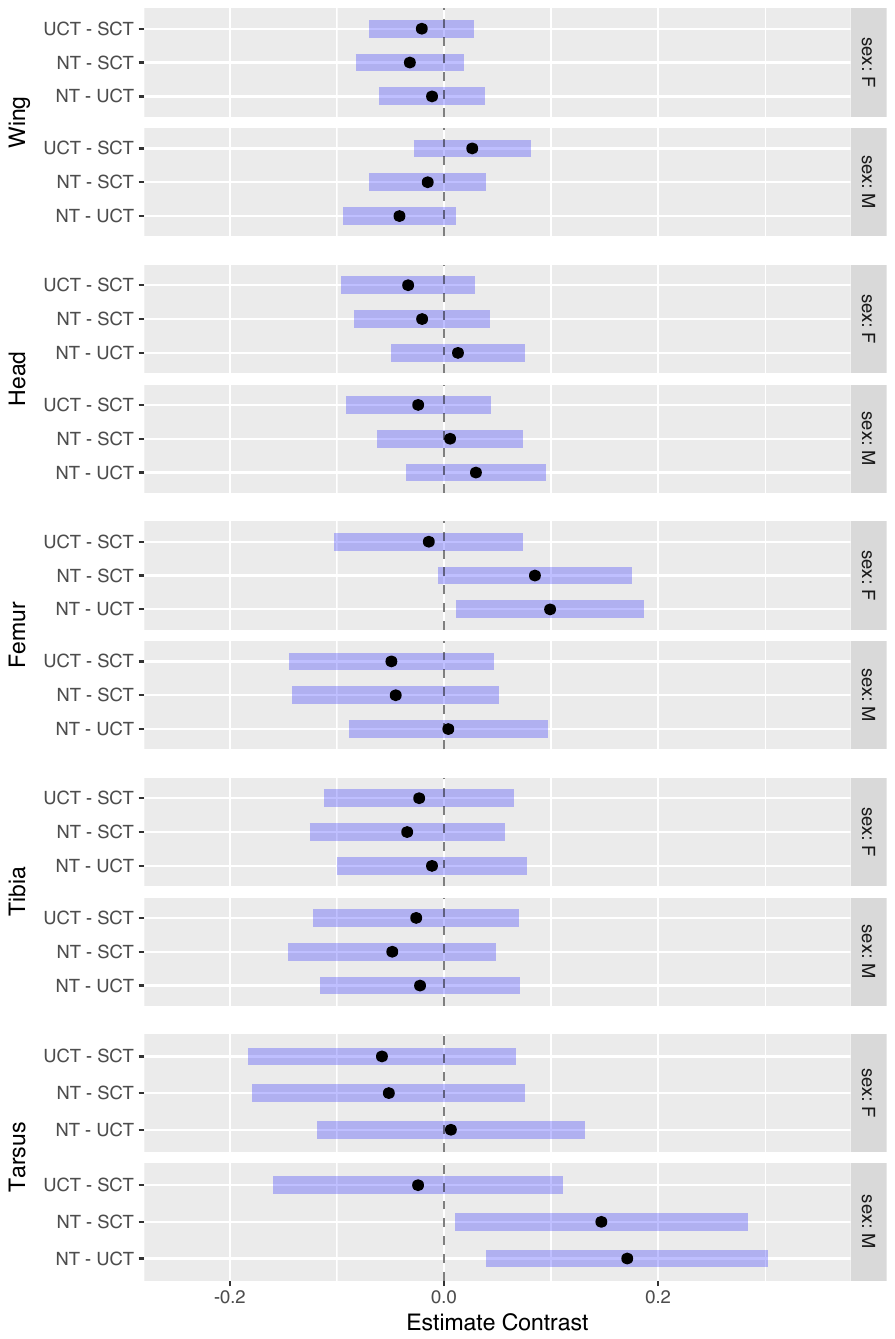

Figure S5: Generation 35 Allometry contrasts, and their 95% confidence intervals from model estimates. Measurements were log_2_ transformed before modelling and contrasts were computed with emmeans.
